# Gene duplication to the Y chromosome in Trinidadian Guppies

**DOI:** 10.1101/2021.02.16.431319

**Authors:** Yuying Lin, Iulia Darolti, Benjamin L. S. Furman, Pedro Almeida, Benjamin A. Sandkam, Felix Breden, Alison E. Wright, Judith E. Mank

## Abstract

Differences in allele frequencies at autosomal genes between males and females in a population can result from two scenarios. Unresolved sexual conflict over survival produces allelic differentiation between the sexes. However, given the substantial mortality costs required to produce allelic differences between males and females at each generation, it remains unclear how many loci within the genome experience significant sexual conflict over survival. Alternatively, recent studies have shown that similarity between autosomal and Y sequence, arising from duplication onto the Y, can create perceived allelic differences, and this represents potentially resolved sexual conflict. However, Y duplications are most likely in species with large non-recombining regions, in part because they simply represent larger targets for duplications. We assessed the genomes of 120 wild-caught guppies, which experience extensive predation- and pathogen-induced mortality and have a relatively small ancestral Y chromosome. We identified seven autosomal genes that show allelic differences between male and female adults. Five of these genes show clear evidence of whole or partial gene duplication to the Y chromosome, suggesting that the male-specific region of the guppy Y chromosome, although relatively small, may nonetheless act as a hotspot for the resolution of sexual conflict. The remaining two genes show evidence of partial homology to the Y. Overall, our findings suggest that the guppy genome experiences a very low level of unresolved sexual conflict over survival, and instead the Y chromosome, despite its small ancestral size and recent origin, acts as a major mechanism of conflict resolution.

## Introduction

Differences in allele frequency between males and females at autosomal loci can result from two alternative sources. Numerous recent studies have used allelic differences between males and females within a population (F_ST_, D_xy_, etc) as a way to infer sex-differences in viability or survival, and therefore sexual conflict over mortality (Cheng & Kirkpatrick, 2016; Dutoit et al., 2018; Flanagan & Jones, 2017; Kasimatis et al., 2021; Lucotte et al., 2016; Wright et al., 2018, 2019).This approach assumes that allele frequencies are the same in males and females at conception, but diverge over the course of a generation for loci with alleles that benefit the survival of one sex at some survival cost to the other (intra-locus sexual conflict). We expect that sexual conflict over survival would produce a signature of allelic differentiation between the sexes as well as balancing selection (Mank, 2017). The latter results from all forms of intra-locus sexual conflict, not just that over survival, as alleles are selected for or against depending on whether they are present in males or females (Barson et al. 2015; Connallon & Clark 2014; Foerster et al. 2007; Hawkes et al. 2016; Johnston et al. 2013; Lonn et al. 2017; Ruzicka et al. 2019). Balancing selection resulting from sexual conflict may be a major factor in the maintenance of genetic polymorphisms within populations (Connallon & Clark 2014), therefore, the proportion of loci subject to unresolved sexual conflict within the genome, as well as the types of loci affected, may have important implications for a range of evolutionary factors, such as the speed and genetic basis of adaptation.

Intra-locus conflict over survival implies a significant mortality cost each generation. Modelling and simulation methods suggest that the sex-specific mortality rates necessary to generate significant allelic differences between the sexes within each generation are quite high for any one locus (Bissegger et al. 2019; Kasimatis, Ralph, & Phillips 2019; Ruzicka et al. 2020). This precludes the presence of large numbers of sites subject to sexual conflict due to survival, as the associated mortality load would simply be too great. Moreover, recent work has highlighted the potential that many perceived allelic sex differences between the sexes actually are the result of sequence homology between autosomal and sex-linked loci (Bissegger et al. 2019; Kasimatis et al. 2019; Mank, Shu, & Wright 2020; Ruzicka et al. 2020). The Y chromosome may preferentially retain duplicates that play an important role in male development or fitness (Bachtrog, 2013; Carvalho et al., 2015), and the process of Y duplication ironically offers a route to sexual conflict resolution even though it may lead to the mistaken perception of unresolved conflict acting on autosomal loci.

It therefore remains unclear how common and pervasive sexual conflict over survival is in natural populations, or what types of loci, if any, it is expected to target. Here we assess the potential for intersexual F_ST_ based on a sample of 120 wild-caught adult guppies, which are expected to experience substantial predation- and pathogen-induced mortality. We identified seven genes that show evidence of allelic differences between adult males and females. Of those seven genes, five show evidence of Y duplicates, and two have sex-specific functions, including germline differentiation and sex hormone signaling. This is surprising, as although the size of the guppy Y varies substantially across populations and related species (Wright et al. 2017; Darolti et al. 2019; Ameida et al. 2021), the conserved, ancestral part of the Y is very small, spanning at most 5 Mb (Almeida et al., 2020; Darolti et al., 2019; Fraser et al., 2020). Given this small size, we might expect that the Y chromosome in guppies does not represent a large enough genomic target for sexual conflict resolution through gene duplication. Instead, our results show that the guppy Y, although a very small region of the genome, may indeed act as a hotspot for the resolution of conflict.

## Results

We individually sequenced 60 male and 60 female wild-caught adults from three rivers in Trinidad to an average of ~30X coverage in males and ~20X coverage in females after filtering, trimming and quality control (see Materials and Methods for further details). The greater read depth in males was designed to aid detection of the Y chromosome, which is present in only one copy in each genome. Because the size of the non-recombining region of the sex chromosome varies across guppy populations (Almeida et al., 2020; Wright et al., 2017), and sex chromosome divergence results in allelic differences on the X and Y between males and females, we confined all our analyses to autosomal sequence, although we present the X chromosome (Chromosome 12) in figures for comparison.

From our sequencing data, we obtained 7,889,657 biallelic high quality filtered autosomal SNPs. We further filtered these to 253,375 SNPs present in annotated autosomal coding sequence. We focused on coding sequence for three reasons. First, non-coding sequence is enriched for repetitive elements, and the accumulation of repetitive elements on the ancestral guppy Y chromosome (Almeida et al., 2020) would lead to male-specific SNPs in repetitive elements that could bias our estimates of Y duplications. Second, previous work has suggested that unresolved sexual conflict on the autosomes occurs primarily in coding sequence (Ruzicka et al., 2019), and this fits with the substantial evidence that sexual conflict over gene regulation can be relatively quickly resolved through sex-specific gene regulation (Kopp, Duncan, & Carroll 2000; Mank 2017; Wright et al. 2019). Finally, by focusing on functional coding sequence, we are able to determine the potential functional role of genes that are subject to sexual conflict.

### Intersexual F_ST_

Given the low level of allelic divergence expected between males and females, it is critical to minimize false positives (Kasimatis et al. 2019). We therefore used three independent methods to estimate SNPs with elevated intersexual F_ST_. We identified SNPs that were 1) in the top 1% of the autosomal F_ST_ distribution 2) were significant after permutation testing of samples (1000 replicates, P < 0.001) and 3) showed significant differences in male and female allele frequency based on Fisher’s exact test (P < 0.001) (Supplementary Fig. 1). We identified 504 autosomal coding sequence SNPs that were significant by all three of these measures (Supplementary Figs. 1 and 2), designated as high intersexual F_ST_ SNPs.

We then identified autosomal coding genes with ≥ 3 high intersexual F_ST_ SNPs (Bissegger et al. 2019; Tobler, Nolte, & Schlötterer 2017), resulting in seven loci with significant sexual differentiation. For each of these genes, average intersexual F_ST_ was significantly greater than the autosomal average (Table 1, Supplementary Fig. 3).

**Table 1.** Summary statistics of sexually differentiated genes. Statistically significant values are shown in bold.

| ENSPREG<sup>1</sup><br>(Gene Name if known) | Chrom | Mean<br>F<sub>ST</sub><sup>2</sup> | M:F read<br>depth<sup>3</sup> | M:F SNP<br>density<sup>3</sup> | Gene functions<sup>4</sup> |
| --- | --- | --- | --- | --- | --- |
| 03921<br>(Olr1496-like) | LG7 | **0.1645** | **1.3809** | 1.3000 | Immune response & germline differentiation |
| 03854<br>(si:rp71-17i16.5) | LG7 | **0.0892** | **1.1580** | 1.1200 | Phosphatidylinositol 3-kinase signalling, involved in immune, inflammatory and allergic responses |
| 09757<br>(BCL11A) | LG15 | **0.0419** | **1.3280** | 1.0000 | Hematopoietic cell differentiation & brain development |
| 07572<br>(Fezf1) | LG6 | **0.0412** | **1.5082** | 1.0077 | Nervous system development, migration of gonadotropin-releasing hormone neurons |
| 05754 | LG15 | **0.0238** | **1.8887** | 0.9445 | N/A |
| 15023 | LG18 | **0.0237** | 1.0829 | 1.2143 | N/A |
| 12439 | LG22 | **0.0098** | 1.0802 | 1.0417 | N/A |

1 Ensembl guppy gene number
2 Significance based on Wilcoxon's rank-sum test ( $P < 0.01$ ), compared to autosomal average.
3 Significance based on Fisher's Exact Test ( $P < 0.05$ ), relative to genome-wide M:F SNP density.
4 From Gene Ontology (Ashburner et al., 2000; "The Gene Ontology resource: enriching a GOld mine," 2021) and Gene Cards (Stelzer et al., 2016).

Estimates of intersexual F_ST_ and Tajima’s D can be biased due to relatedness of individuals within groups. Although our sampling design was balanced, with 10 females and 10 males collected at two separate sites for each of three rivers across the island of Trinidad (see Almeida et al. 2020 for details), we could not account for relatedness in wild-caught samples at the time of collection. We therefore assessed pairwise kinship coefficients among all our samples using both KING (Manichaikul et al., 2010) and NgsRelate (Korneliussen & Moltke 2015), as implemented in ANGSD (Korneliussen, Albrechtsen, & Nielsen 2014). Neither method identified greater relatedness among male compared to female samples (Supplementary Fig. 4).

### Y duplications

When mapping whole-genome data to a female reference genome, sequence similarity between the male-specific Y chromosome and the autosomes can lead to the perception of allelic differences between males and females. Recent work has shown that many genes with allelic sex differences are in fact autosomal loci that either have recent Y duplicates (Bissegger et al. 2019; Mank et al. 2020) or otherwise display sequence homology to the Y (Kasimatis et al., 2021). We can use differences in M:F read depth to identify these genes (Hall et al., 2013). Duplication of the complete coding sequence for an autosomal gene on the Y chromosome would produce an average M:F read depth of 1.5 (three copies in males, two in females), and subsequent tandem duplications on the Y would result in M:F read depth > 1.5. Partial Y duplications, or a full duplication followed by significant differentiation, would result in average M:F read depth > 1 and < 1.5 when mapping to a female genome after correcting for differences in library size.

Five of our seven sexually differentiated genes showed average M:F read depth significantly > 1 (Table 1, Supplemental Fig. 5) consistent with at least a partial duplication of the coding sequence on the Y chromosome (Fig. 1). Four of these Y-duplicated genes have annotated functions, two of which (Olr1496-like and Fezf1) have a known role in sexual differentiation (Table 1), as is expected for genes on the Y chromosome.

**Fig. 1.**
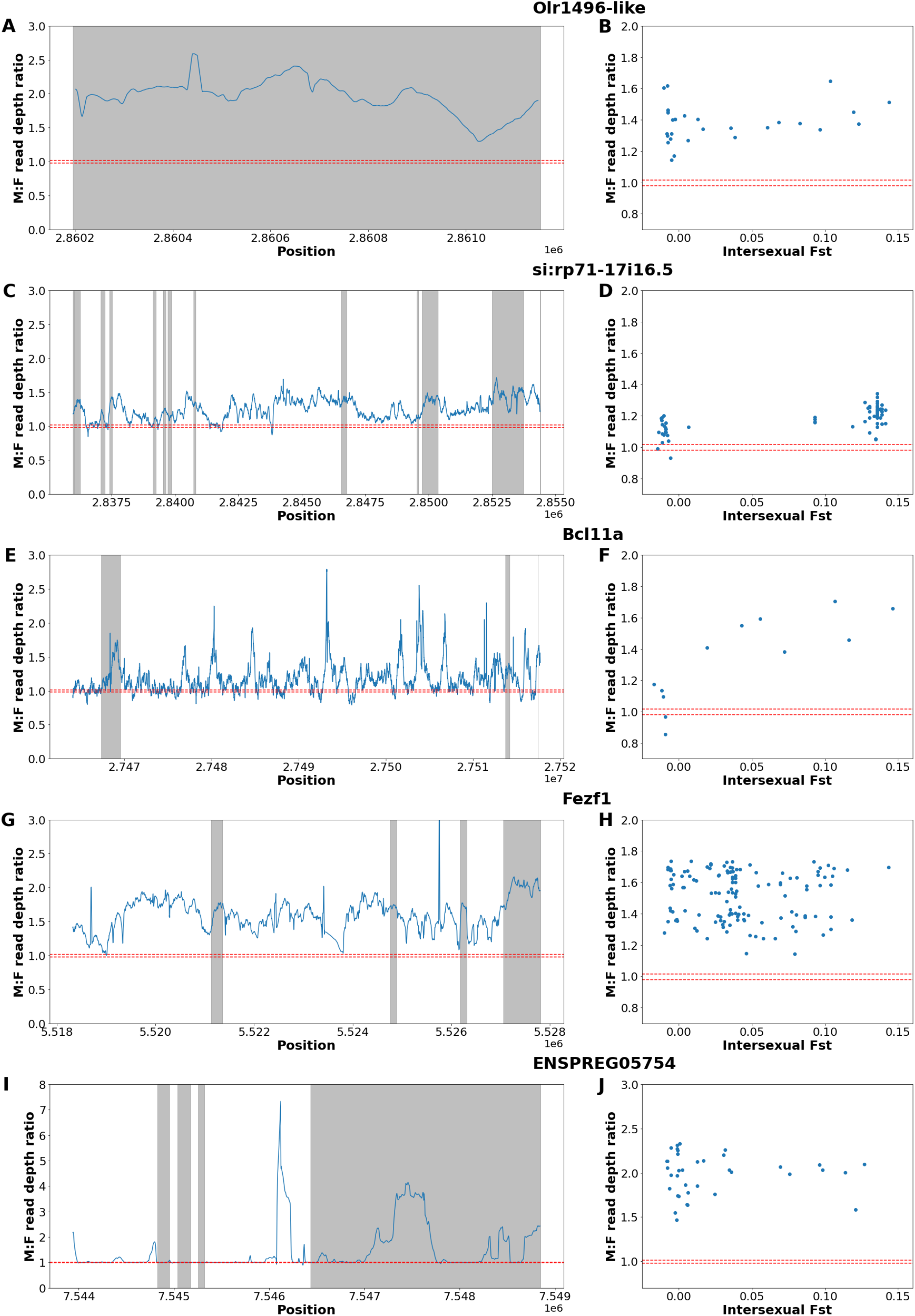
Elevated M:F read depth ratio indicates partial gene duplication of Olr1496-like (Panels A and B), si:rp71-17i16.5 (Panels C and D), BCL11A (Panels E and F), Fezf1 (Panels G and H), ENSPREG05754 (Panels I and J). Left panels show M:F read depth as a function of physical position (in Mb), with coding regions shaded in grey. Panels on the right show intersexual F_ST_ and M:F read depth ratio for each SNP. 95% autosomal confidence intervals for M:F read depth are indicated by horizontal red lines.

In order to differentiate whether the pattern of elevated M:F F_ST_ is due to limited sequence homology between the autosomal genes and the Y chromosome, or results from at least partial gene duplication, we mapped the pattern of M:F read depth across each of these 5 genes (Fig. 1). The high M:F read depth for Olr1492-like, consistent across the full length of the single exon of this gene, suggests a complete duplication, possibly in two copies (Fig. 1A). Similarly, the M:F read depth ratio for si:rp71-17i16.5 (Fig. 1C), Bcl11a (Fig. 1E), and Fezf1 (Fig. 1G) suggests a single or double copy duplication, of just the exons, possibly through retrotransposition. The high M:F read depth in ENSPRG05754 (Fig. 1I) suggests a partial Y duplication of the last exon of the gene, likely present in several Y copies.

Given sufficient evolutionary time, Y duplications will also accumulate male-specific SNPs (Tobler et al. 2017), leading to a signal of elevated M:F SNP density. Of the five sexually differentiated genes with greater read depth in males, none showed significantly higher M:F SNP density (Table 1), although all values were >1, suggesting that duplication to the Y is relatively recent, and consistent with the recent origin of this chromosome (Darolti et al. 2019). Across our seven sexually differentiated genes, we did not observe evidence of duplications to the X chromosome, which would result in three copies in males and four in females and therefore an average M:F read depth significantly <1.

### Sexual conflict over survival for autosomal genes

We identified just two sexually differentiated genes without significantly higher M:F read depth (Table 1), both of which displayed relatively modest levels of intersexual F_ST_ (0.0237 and 0.0098). At first consideration, this small number of loci with modest F_ST_ is consistent with limited unresolved sexual conflict and the expectation that large numbers of high intersexual F_ST_ loci would require unsustainable levels of sex-specific mortality each generation (Bissegger et al. 2019; Kasimatis et al. 2019; Ruzicka et al. 2020).

Genes subject to ongoing sexual conflict are expected to experience balancing selection, which is often measured with Tajima’s D (Bissegger et al. 2019; Cheng & Kirkpatrick 2016; Mank 2017; Wright et al. 2018). We therefore compared Tajima’s D for these two genes to the autosomal average, genes with known immune function, which are known to be subject to high levels of balancing selection (Andrés et al. 2009; Ferrer-Admetlla et al. 2008; Van Oosterhout 2009; Weedall & Conway 2010), and to other genomic categories (Fig. 2). Contrary to our expectation of elevated Tajima’s D for these two sexually differentiated genes, neither showed a Tajima’s D value significantly greater than the autosomal average, and Tajima’s D for ENSPREG15023 is significantly lower than the autosomal average (Fig. 2).

**Fig. 2.**
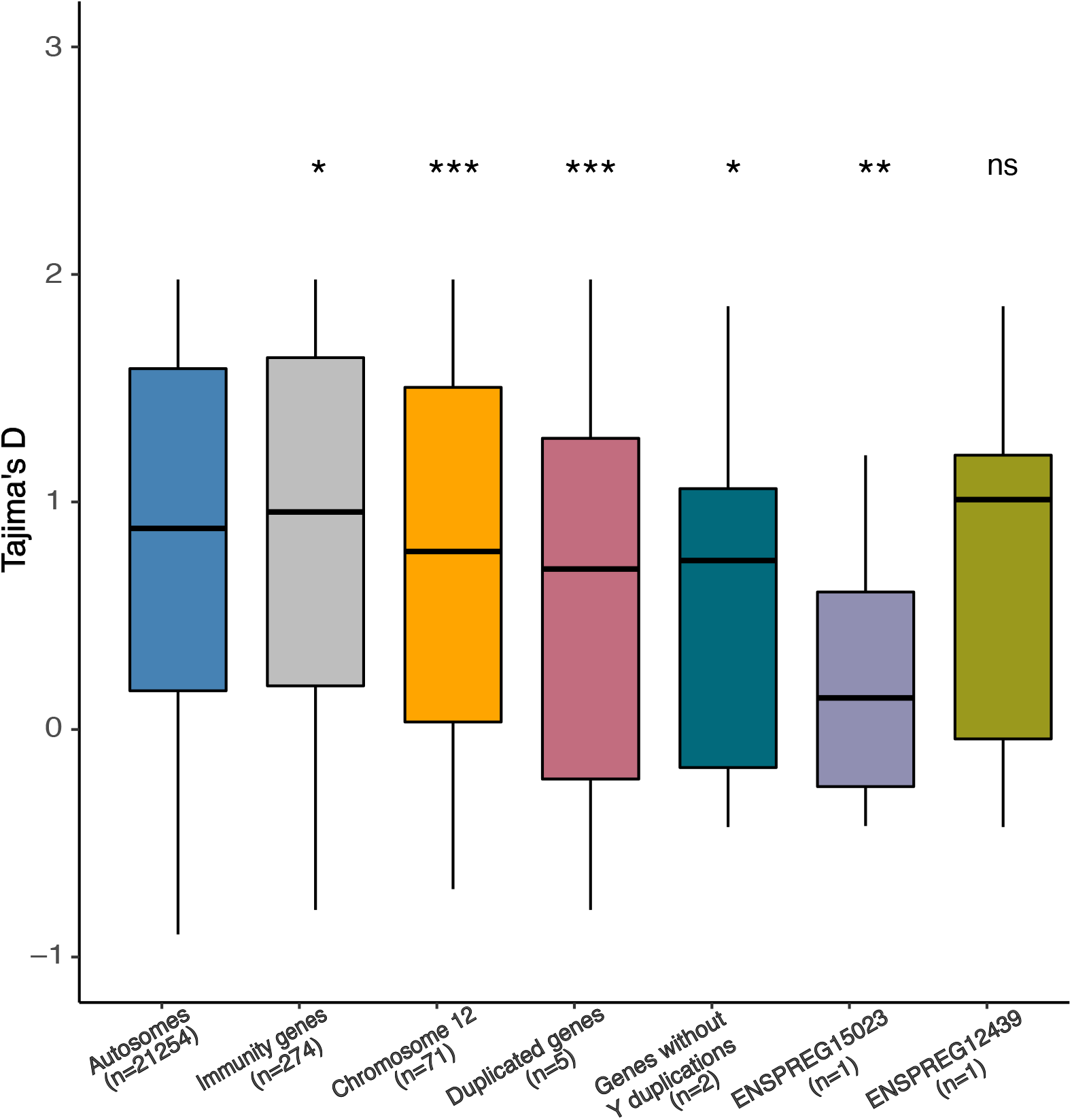
Distribution of Tajima’s D among different gene categories. Significant differences to autosomal distribution are indicated (Wilcoxon rank-sum test, *P < 0.01, **P < 0.001, ***P < 0.0001, ns: non-significant).

Many things can influence Tajima’s D estimates, and the lack of an elevated signature of balancing selection relative to the remainder of the genome is not necessarily indicative of a lack of sexual conflict. In order to understand the dynamics of these two genes in more detail, we mapped M:F read depth as a function of genomic location and intersexual F_ST_ (Fig. 3). Indeed, we found evidence of elevated M:F read depth for ENSPREG15023 across most of the exon, consistent with either partial duplication or full exonic duplication to the Y followed by partial degeneration. Many of the high F_ST_ SNPs in the gene have a high M:F read depth, consistent with Y duplication or significant Y homology. The pattern for ENSPREG12439 is more consistent with significant Y homology given the highly variable M:F read depth across the gene. Importantly, all SNPs with high M:F F_ST_ values for this gene also show high M:F read depth. These results, coupled with the lack of elevated Tajima’s D, suggest that these two genes also exhibit significant homology to the Y, and that this is driving the elevated intersexual F_ST_ instead of unresolved sexual conflict over survival.

**Fig. 3.**
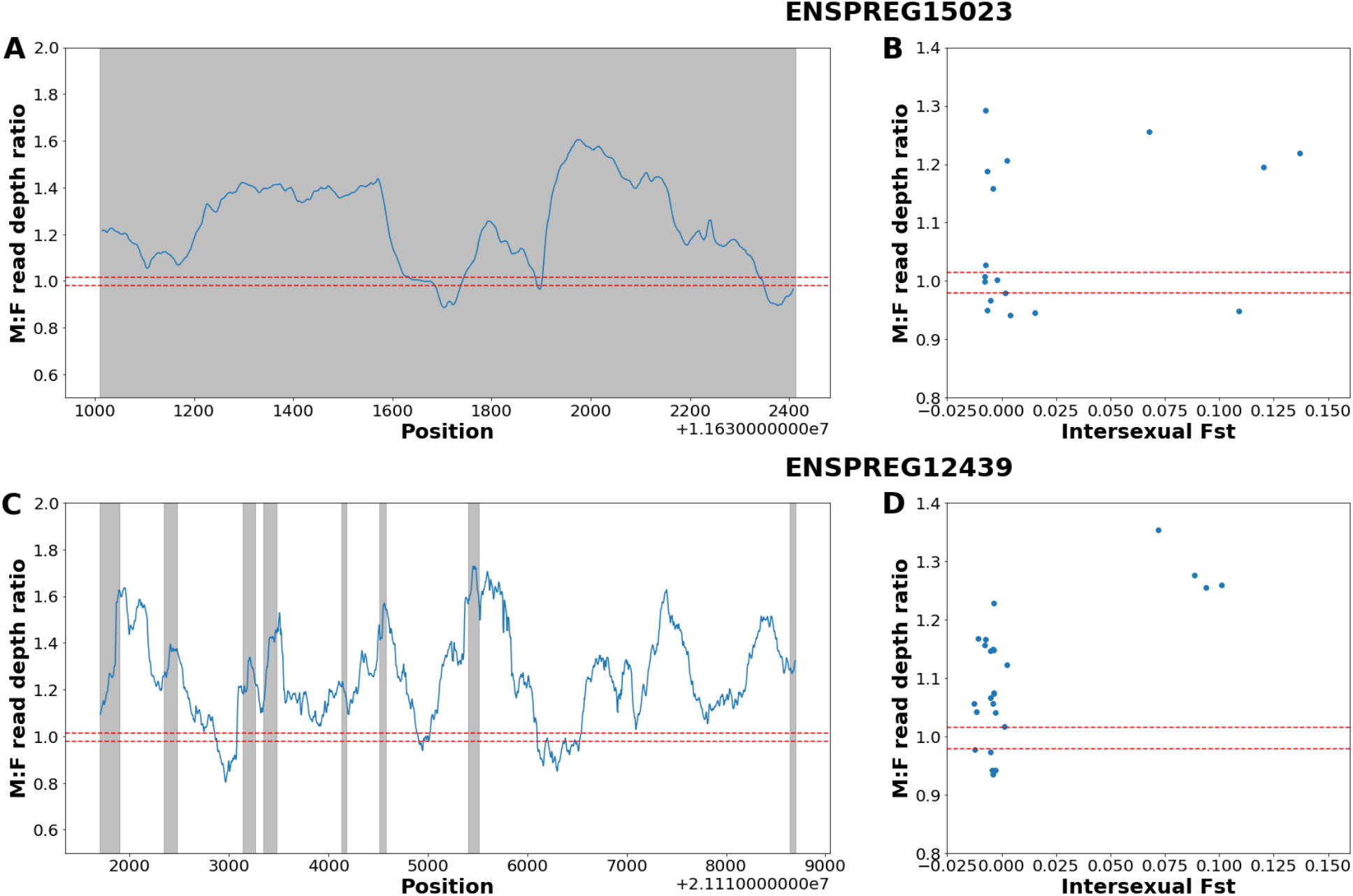
Normalized M:F read depth ratio for ENSPREG15023 (panels A and B) and ENSPREG12439 (panels C and D). Left panels show M:F read depth as a function of physical position (in Mb), with coding regions shaded in grey. Panels on the right show intersexual F_ST_ and M:F read depth ratio for each SNP. 95% autosomal confidence intervals for M:F read depth are indicated by horizontal red lines.

## Discussion

We identified 504 coding sequence SNPs in the guppy autosomal genome which showed significant differences in allele frequency between males and females. We used these to identify seven autosomal genes with significant average intersexual F_ST_. Our approach, based on the intersection of three statistical methods, reduces the likelihood of false positives and results in a high confidence gene list of intersexual F_ST_ (Table 1, Supplementary Figs. 1, 2, and 3). This list of genes can be used to assess the relative role of Y duplication in resolving conflict and driving intersexual F_ST_, versus the role of sex differences in mortality in sexual conflict of autosomal genes.

### The Y chromosome in guppies as a locus for conflict resolution

Five of the seven high-confidence sexually differentiated genes showed evidence of Y duplications based on elevated male-to-female read depth ratios (Table 1, Supplementary Fig. 4). Two of these (Olr1496-like and Fezf1) have sex-specific functions based on Gene Ontology and Gene Cards annotations, consistent with theory and empirical studies showing that Y chromosomes can accumulate gene duplicates with male-specific functions (Carvalho et al. 2015; Mahajan & Bachtrog 2017). Specifically, Olr1496 (Olfactory factor 1496-like gene) plays a role in multiple aspects of germline development in *C. elegans* (Cho, Rogers, & Fay 2007), and Fezf1 (FEZ Family Zink Finger 1) plays an important role in the migration of gonadotropin-releasing neurons (Damla Kotan et al., 2014).

Y chromosomes in general represent a unique genomic environment. Although Y chromosome gene content degrades quickly following recombination suppression (Bachtrog, 2013), several recent studies indicate that older, large Y chromosomes, such as that in *Drosophila* (Carvalho et al. 2015; Koerich et al. 2008; Tobler et al. 2017), sticklebacks (Bissegger et al. 2019) and humans (Kasimatis et al. 2020), contain substantial numbers of genes duplicated from autosomes. Duplications may represent an important mechanism of sex chromosome divergence, and the Y chromosome may preferentially retain duplicates that play an important role in male development or fitness (Bachtrog, 2013; Carvalho et al., 2015), offering a hotspot for sexual conflict resolution within the genome.

However, it remains unclear how common Y duplications of autosomal genes occur in younger systems with a far smaller male-specific region. Although the size varies by population, the conserved non-recombining region of the Y chromosome in guppies is relatively small, spanning at most 5 Mb (Almeida et al., 2020; Darolti et al., 2019). Others have failed to recover evidence for even this limited region of Y degeneration (Charlesworth et al., 2020; Fraser et al., 2020). Our findings add further support for a region of recombination suppression across populations on Trinidad, as only a male-specific region of the Y can explain the M:F read depth differences we observe. Moreover, our work suggests that the guppy Y chromosome is dynamic with regard to gene content, and acts as a hotspot for gene duplications with male-specific functions despite its recent origin and small size. It is also possible that Y genes have duplicated to the autosomes, which would also produce a pattern of increased male read depth, although this is arguably less likely. Taken together, our work suggests that even homomorphic sex chromosomes may act as a hotspot of sexual conflict resolution. Moreover, our results further emphasize the importance of accounting for Y gene duplications in scans for M:F F_ST_, as the all of our sexually differentiated genes show evidence of Y duplication.

The Y-duplicated genes we identify here are not present in our previous list of Y-linked genes based on male-specific sequence (Almeida et al., 2020), or in other similar analyses (Fraser et al., 2020). This is not surprising, as duplications, particularly if recent, will still retain substantial homology to the autosomal copy and will not be detected when bioinformatically identifying sequence that is unique to male genomes. Consistent with recent duplications and limited divergence, none of our Y-duplicated genes exhibit significantly elevated average M:F SNP density (Table 1, Fig. 2), although the values are all >1.

We have previously (Wright et al., 2018) noted evidence of intersexual F_ST_ in guppies (*Poecilia reticulata*). This finding was curious given that the data from this earlier work derived from a lab population, free of most of the pathogens and all the predators that would exacerbate sex-differences in mortality and predation (Wright et al., 2018). It was not clear how much of this signal, if any, was due to Y duplications. However, it is worth noting that elevated intersexual F_ST_ was highest for genes with male-biased expression, as would be expected for genes with Y duplicates. We also did not observe a concomitant pattern of elevated Tajima’s D for these genes, which is inconsistent with sexual conflict over survival.

### Sexual conflict over survival targets few genes in the guppy autosomal genome

Intra-locus sexual selection over survival or viability leads to allele frequency differences between the sexes over the course of a generation, as an allele increases the survival of one sex at a mortality cost to the other (Kasimatis et al. 2019; Mank 2017; Wright et al. 2018). The significant mortality costs required each generation to generate allele frequency differences between the sexes preclude large numbers of genes with significant M:F F_ST_ (Bissegger et al. 2019; Kasimatis et al. 2019; Ruzicka et al. 2020). At most, we might expect a limited number of loci with significant allelic differentiation between males and females, and this would be most evident in wild species where males and females experience differences in predation, parasite or pathogen loads. Despite the fact that we sampled wild individuals of a classic prey species with very high potential for sexual conflict over survival, we observe just two loci in the genome that show evidence of sexual conflict over survival based on M:F F_ST_. Both of these loci exhibit very low M:F F_ST_.

We expect that if these genes are indeed subject to sexual conflict over mortality, we would observe elevated Tajima’s D (Fig. 2), however this was not the case for either locus. Moreover, both of these loci exhibit patterns of M:F read depth consistent with significant Y homology (Fig. 3). This suggests that the potential for sexual conflict over survival is quite low, even in a species where we most expect to observe it.

### Concluding remarks

Here we use a high-stringency filtering method to detect genes within the guppy genome that exhibit population genetic signatures expected from sexual conflict over survival. Although wild guppies are expected to have high potential for sexual conflict over survival, we in fact found no genes within the genome with patterns consistent with this. Instead, despite the small size of the conserved non-recombining region of the guppy Y, we observe five loci that show patterns consistent with autosome-to-Y duplications, and two more that are suggestive of Y homology of without duplication. This highlights the potential of even young, small Y chromosomes as regions of conflict resolution.

## Materials and Methods

### Data Collection and Genotyping

Samples were collected from three rivers, Aripo, Yarra, and Quare, in Trinidad in December 2016, in accordance with national collecting guidelines. In total, 10 males and 10 females were collected from one high predation and one low predation population in each river, resulting in 120 samples, which were sequenced individually with Illumina HISEQX. Further sequencing details are available in Almeida et al. (2020).

We used FastQC v0.11 (http://www.bioinformatics.babraham.ac.uk/projects/fastqc) and Trimmomatic 0.36 (Bolger et al., 2014) to remove adapter sequences and low-quality reads.

After quality control, we recovered ~30X average sequencing depth for males and ~20X sequencing depth for females. High quality reads were aligned against the *Poecilia reticulata* female reference genome (Ensembl GCA_000633615.2) (Künstner et al., 2016), using BWA 0.7.15 MEM algorithm (Li & Durbin 2009) with default parameters. We filled in mate coordinates and mate related flags, sorted alignment by coordinates, and marked PCR duplications with SAMtools-1.9 (Li et al., 2009).

We called genotypes across all the samples using the ‘mpileup’ function from bcftools with the following parameters: --min-MQ 20 --min-BQ 20 --skip-indels -a FORMAT/AD, FORMAT/DP. After genotyping, we used VCFtools v0.1.16 (Danecek et al., 2011) to exclude SNPs that had either: (1) genotype quality < 25; (2) sequencing depth <0.5x or >10x of average depth; (3) missing data in > 10% of individuals; (4) minor allele frequency < 0.05 or (5) relative read depth between reference allele and alternative allele < 0.2 or > 0.8. In total, the autosomal filtered SNP dataset consisted of 7,889,657 biallelic SNPs. We extracted 253,375 SNPs in annotated coding sequences (Ensembl build Guppy Female 1.0) for downstream analysis. Finally, we confined our analysis to autosomal genes, but included the X chromosome (Chromosome 12) in figures as a means of comparison.

### Intersexual F_ST_

In order to estimate intersexual allele frequency differences, we implemented Weir & Cockerham’s estimator of F_ST_ (Weir & Cockerham 1984) between males and females using VCFtools v0.1.16 for each SNP in genome-wide coding sequence regions. We employed three methods jointly to identify SNPs exhibiting high F_ST_. First, we used a cut-off method, retaining SNPs in only the top 1% of autosomal F_ST_ values. Second, we performed permutation tests by randomly assigning individuals to one of two sex groups to generate a null distribution of F_ST_ across the genome. We determined significance for each SNP from 1000 replicates, using a P < 0.001 threshold. Finally, we performed Fisher’s exact test on SNPs to determine significance of allele frequency differences between males and females (P < 0.001). We denoted SNPs that were significant in all three of these measures as high F_ST_ SNPs.

Using approaches to further limit false positives (Bissegger et al., 2019; Tobler et al., 2017), we identified 7 genes with ≥ 3 high intersexual F_ST_ SNPs, which we designated as sexually differentiated genes. We calculated average intersexual F_ST_ for all genes using VCFtools v0.1.16, respectively. We used Wilcoxon rank-sum test to indicate statistical difference in intersexual F_ST_ between autosomal genes and other gene categories (sexually differentiated genes and genes on the sex chromosome).

### Relatedness Inference

In order to avoid biases in calculating intersexual allele frequency differences due to relatedness among individuals, we used KING 2.2.7 (Manichaikul et al., 2010) to infer the pairwise degree of relatedness between individuals from estimated kinship coefficients. We first converted genotype data from the raw, unfiltered SNPs dataset to plink binary format using PLINK 1.9 (Purcell et al., 2007). In order to avoid potential biases from KING software (Ramstetter et al., 2017) and validations, we also used NgsRelate v2 (Korneliussen & Moltke 2015), implemented in ANGSD (Korneliussen et al. 2014) to infer genetic relatedness coefficients for each pair of individuals.

### Assessing Y Duplications of Sexually Differentiated Genes

When using a female reference genome, reads from genes which have duplicated to the male-specific region of the Y chromosome will map back to original autosomal or X chromosome regions, resulting in elevated M:F coverage (Bissegger et al. 2019; Mank et al. 2020). For example, if an autosomal gene has one Y duplication, we would expect three copies in males (two autosomal and one Y-linked) and two copies in females, and therefore an average M:F read depth of 1.5. We first calculated M:F read depth for each coding sequence SNP from genotype data, as male coverage/female coverage, correcting for differences in average genomic read depth between males and females. In order to validate the M:F read depth ratio of sexually differentiated genes, we calculated male and female normalized read depth for 7 genes for each base pair, based on male and female pooled reads.

Genes on the Y chromosome will accumulate male-specific mutations over time, leading to an increased number of male-specific mutations as well as elevated M:F SNP density (Bissegger et al. 2019; Mank et al. 2020). M:F SNP density for each gene was calculated as number of male SNPs/number of female SNPs.

### Tajima’s D

Based on the filtered genotype data, we calculated Tajima’s D for the coding sequence of all autosomal genes using VCFtools v0.1.16. We compared mean Tajima’s D for autosomal genes (excluding those with immune function), genes with immune function (defined following Wright et al. 2017, Wright et al. 2018), and our seven sexually differentiated genes.

### Functional Annotations

The small number of sexually differentiated genes precludes Gene Ontology enrichment analysis. We therefore cataloged functional annotations from the *Danio rerio* Gene Ontology (Ashburner et al., 2000; “The Gene Ontology resource: enriching a GOld mine,” 2021) and Gene Cards (Stelzer et al., 2016).

## Acknowledgements

This was funded by a Chinese Scholarship Council Doctoral Scholarship to YL (No. 201906040216), a grant from the European Research Council (grant agreement 680951) and a Canada 150 Research Chair to JEM. We thank The Trinidad Ministry of Agriculture, Land and Fisheries and Indar Ramnarine for field permits. We thank XXX for helpful comments and suggestions on the manuscript.

## Data Accessibility Statement

DNA-seq data are publicly available in the NCBI SRA (BioProject ID PRJEB39998). Scripts of all the analysis are available on GitHub XXXXX.

## Author Contributions

J.E.M. and Y.L. designed the study. B.S., F.B. and J.E.M. collected samples. P.A. collected raw sequencing data. Y.L., I.D., B.L.S.F., P.A., B.S., A.E.W. and J.E.M. performed the data analysis. J.E.M., Y.L., I.D., B.L.S.F., B.S., F.B. and A.E.W. contributed to the writing of the manuscript.

---

**Supplementary Fig. 5.**
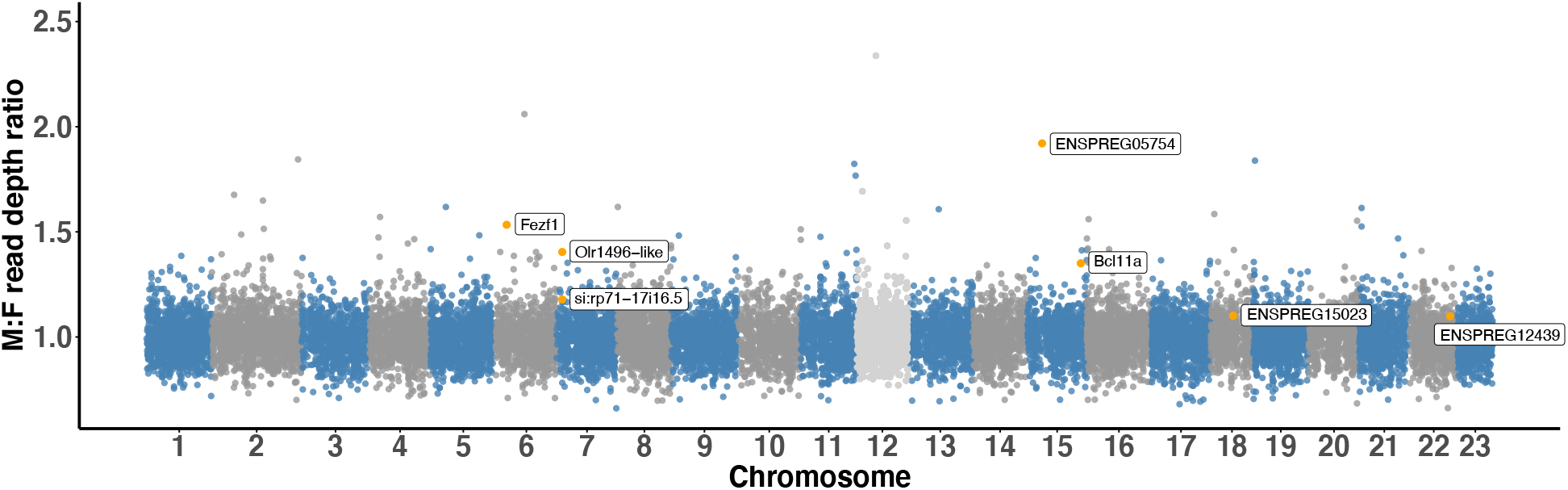
Normalized M:F read depth ratio, with seven sexually differentiated genes indicated in orange. Steel blue and grey: autosomes; Light grey: sex chromosome (Chromosome 12).

## Notes

### Competing Interest Statement

The authors have declared no competing interest.

