## Supplementary Figure 1,2,3,4,5 for "Gene duplication to the Y chromosome in Trinidadian Guppies"

**Supplemental Information**

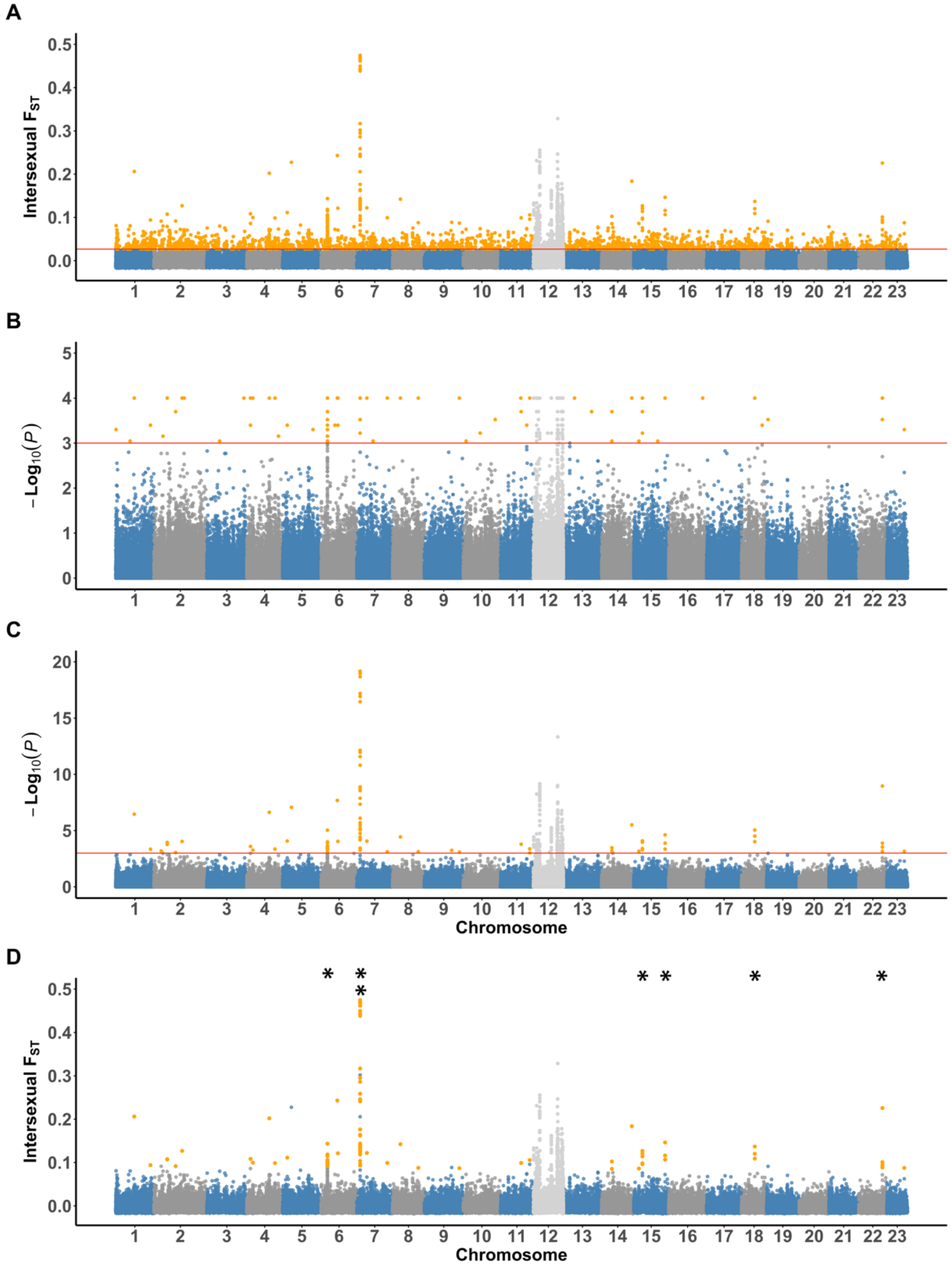


**Supplementary Fig. 1.** Detecting intersexual $F_{ST}$ outliers, using (A) 1% cut-off method, (B) 1000 times permutation test of intersexual $F_{ST}$ ($P < 0.001$) and (C) Fisher's exact test on allele frequency differences ($P < 0.001$). (D) High Intersexual $F_{ST}$ SNPs, based on intersection of A, B and C. * Stars on the top show position of significantly sexually differentiated genes (with at least 3 sexually differentiated SNPs). X-linked genes were excluded from analyses, although we present them here for comparison. Red line: significance level; Steel blue and grey: autosomes; Light grey: sex chromosome (Chromosome 12); Orange: $F_{ST}$ outliers detected.

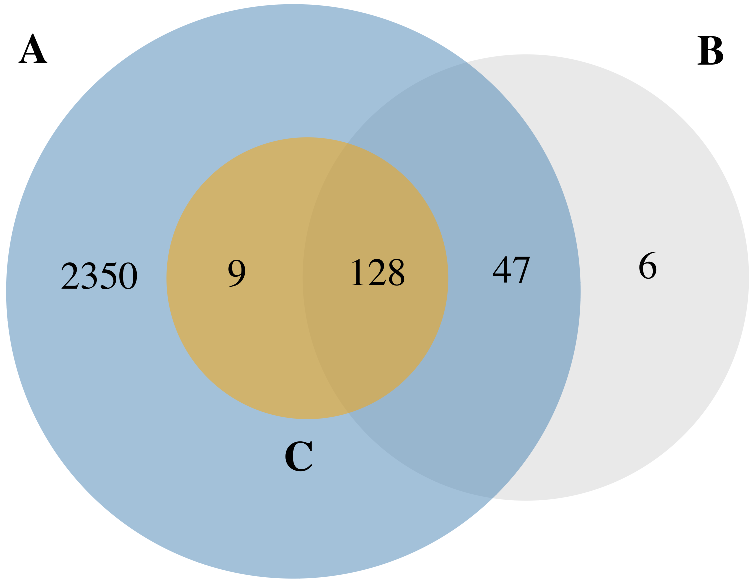


**Supplementary Fig. 2.** Venn diagram showing the number of $F_{ST}$ outliers detected by the (A) 1% cut-off method, (B) permutation test (P < 0.001), (C) Fisher's exact test (P < 0.001) on allele frequency differences. We observe 128 SNPs with elevated intersexual $F_{ST}$ across all three different methods (shown in Supplementary Fig 1D).

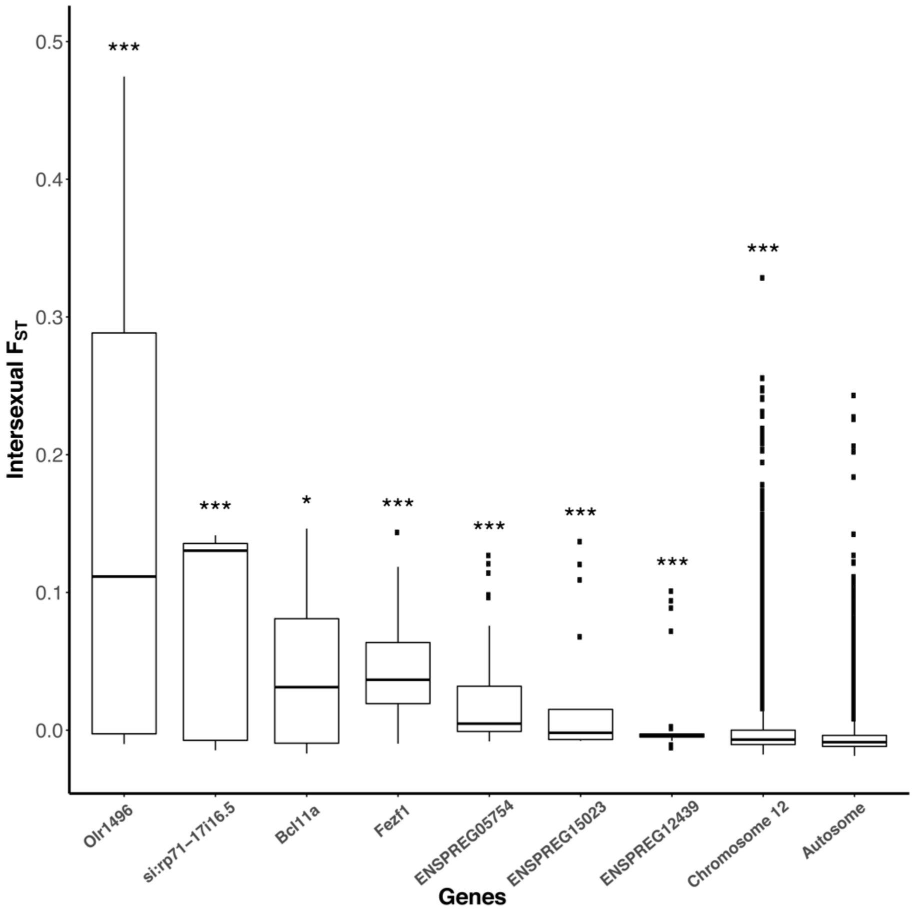


**Supplementary Fig. 3.** Average Intersexual $F_{ST}$ comparison across the 15 sexually differentiated genes, compared to the average on the sex chromosome (Chromosome 12) and the autosomes. Stars indicate significantly elevated intersexual $F_{ST}$ relative to autosomal genes using Wilcoxon rank-sum test. *P < 0.05, **P < 0.01, ***P < 0.001

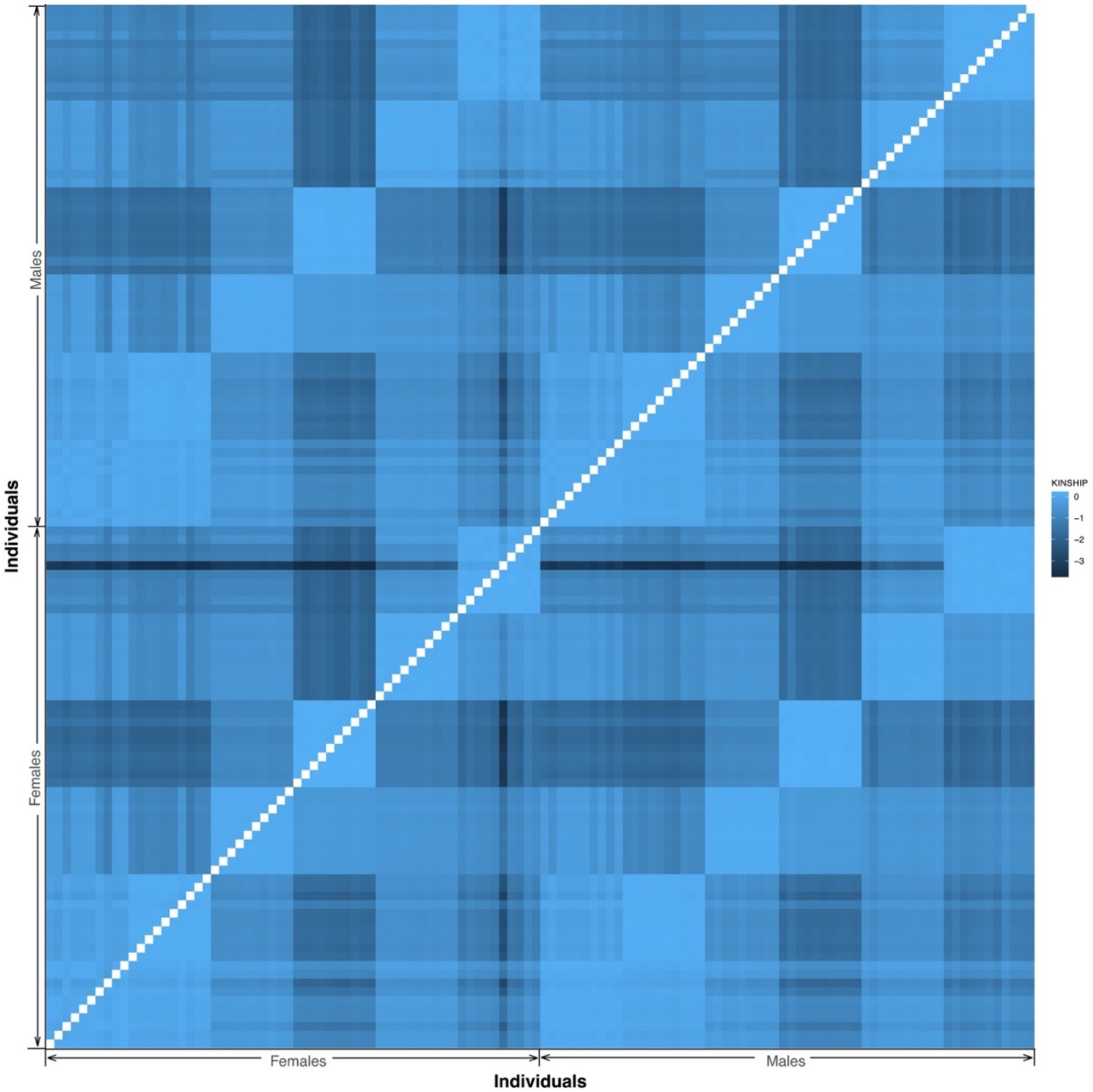


**Supplementary Fig. 4.** Heatmap indicating low pair-wise relatedness between samples in this study. Each column and each row represent one sample. The upper left section shows relatedness inference results from KING; the lower right section shows results from NgsRelate. Intersexual relatedness does not differ from average relatedness across all samples.

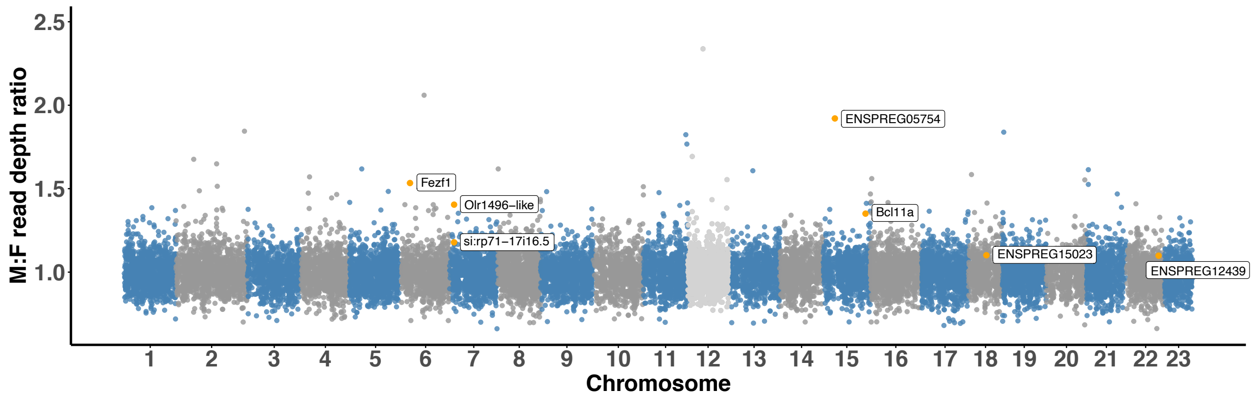


**Supplementary Fig. 5.** Normalized M:F read depth ratio, with seven sexually differentiated genes indicated in orange. Steel blue and grey: autosomes; Light grey: sex chromosome (Chromosome 12).
